# Heterogeneity of Incipient Atrophy Patterns in Parkinson’s Disease

**DOI:** 10.1101/466086

**Authors:** Pedro D. Maia, Sneha Pandya, Justin Torok, Ajay Gupta, Yashar Zeighami, Ashish Raj

**Affiliations:** Department of Radiology, Weill Cornell Medical College of Cornell University 407 E. 61 Street, RR106, New York, NY 10065, USA; Montreal Neurological Institute, McGill University, Montreal, Canada; Department of Radiology, Department of Radiology and Biomedical Imaging, School of Medicine, University of California at San Francisco

**Keywords:** Parkinson’s Disease (PD), brain atrophy, neuropathology, tractography, seed-inference methods, network diffusion models

## Abstract

Parkinson’s Disease (PD) is a the second most common neurodegenerative disorder after Alzheimer’s disease and is characterized by cell death in the amygdala and in substructures of the basal ganglia such as the substantia nigra. Since neuronal loss in PD leads to measurable atrophy patterns in the brain, there is clinical value in understanding where exactly the pathology emerges in each patient and how incipient atrophy relates to the future spread of disease. A recent seed-inference algorithm combining an established network-diffusion model with an L1-penalized optimization routine led to new insights regarding the non-stereotypical origins of Alzheimer’s pathologies across individual subjects. Here, we leverage the same technique to PD patients, demonstrating that the high variability in their atrophy patterns also translates into heterogeneous seed locations. Our individualized seeds are significantly more predictive of future atrophy than a single seed placed at the substantia nigra or the amygdala. We also found a clear distinction in seeding patterns between two PD subgroups – one characterized by predominant involvement of brainstem and ventral nuclei, and the other by more widespread frontal and striatal cortices. This might be indicative of two distinct etiological mechanisms operative in PD. Ultimately, our methods demonstrate that the early stages of the disease may exhibit incipient atrophy patterns that are more complex and variable than generally appreciated.

## 1. Introduction

Parkinson’s Disease (PD) is an age-related progressive neurodegenerative disorder clinically marked by tremor, bradykinesia, and rigidity. Patients exhibit cognitive deficits and with progression develop dementia, depression and/or other neuropsychiatric symptoms (Poewe and Wenning, 2006). Pathologically, these symptoms are caused by the loss of dopaminergic projections in the midbrain from the substantia nigra pars compacta (SN) to the striatum. A common histopathological hallmark of PD is the aggregation of misfolded α-synuclein (AS) protein called Lewy bodies (Jellinger, 2003). Histopathological studies of PD patient brains have strongly suggested that Lewy pathology spreads in a stereotypical stage-wise pattern from the brainstem to subcortical areas and then to the cerebral hemispheres along neuronal pathways (Braak *et al.*, 2003; Del Tredici and Braak, 2016). Recent advances in neuroimaging techniques have confirmed the Lewy pathology staging scheme. Regional MRI-derived atrophy was found to be the highest in SN, followed by atrophy of the striatum, amygdala, hippocampus and other limbic areas. Neocortical atrophy is seen in advanced stages, especially in orbitofrontal, insular and temporo-occipital cortices, in the same general order as seen in pathology stages (Ramírez-Ruiz *et al.*, 2005; Yau *et al.*, 2018).

Despite these advances, we lack understanding of pre-symptomatic stages of PD and it is unclear whether the pathological process begins in the brain or elsewhere in the nervous system. The purpose of this paper is to infer the likely sites of initiation of PD pathology from *in vivo* neuroimaging data, and to understand etiological origins and the sources of heterogeneity amongst PD populations. The basis of our approach is that the spread of PD pathology can be captured deterministically in a network-based model, which can then be “inverted” to enable seed region inference. Here we briefly survey the neuropathological basis of this model and the seed-inference algorithm.

### PD pathology progression occurs on the connectivity network

Toxic misfolded AS undergo template-driven aggregation in a “prion-like” fashion (Wood *et al.*, 1999; Yonetani *et al.*, 2009; Sacino *et al.*, 2013) intracellularly, followed by cell-to-cell transmission via transneuronal pathways along axonal projections to remote areas (Luk *et al.*, 2012; Masuda-Suzukake *et al.*, 2013; Rey *et al.*, 2016). A study showing a wild-type mice receiving a single intrastriatal injection of synthetic AS fibrils led to the trans-neuronal transmission of pathologic AS and Parkinson’s-like Lewy pathology in anatomically interconnected regions (Luk *et al.*, 2012). Lewy pathology spread occurs along the local and long-range fiber projections, thereby suggesting a process of “network spread”. This process is ubiquitous to neurodegenerative diseases, including Alzheimer’s, frontotemporal dementia, and others (Braak and Braak, 1991; Pandya *et al.*, 2017).

Recently, an *in vivo* quantitative approach using predictive network diffusion model (NDM) has been applied to AD and other dementias by specifically modeling the trans-neuronal spread using connectivity (Raj *et al.*, 2012, 2015). The diagnostic and clinical role of NDM and other models of neurodegenerative pathologies through trans-neuronal diffusion is detailed in (Iturria-Medina, 2013; Iturria-Medina *et al.*, 2014; Carbonell *et al.*, 2018). Likewise, a network-based analysis has been applied to a large group of PD patients showing atrophy distribution by modeling the trans-neuronal spread of AS outward from SN (Zeighami *et al.*, 2015). The study was statistically validated using neuropathological and neuroimaging data of 232 PD patients which demonstrated the spatial and temporal atrophy patterns in PD reflecting out of the disease epicenter in the SN. The NDM was validated on the same data series in our laboratory, where we showed that SN-seeded NDM was able to faithfully recapitulate PD cross-sectional atrophy patterns, both at the group level and at the individual level; the latter at lower predictive power and higher variability (Pandya *et al.*, 2018), providing a new computationally-defined staging scheme that mirrors Braak’s six-stage Lewy pathology staging scheme (Braak *et al.*, 2003).

### Utilizing the Network Diffusion model for seed inference

In this paper we report that the NDM can be successfully “inverted” in order to infer the most likely pattern of incipient pathology seeding from available regional atrophy data in PD patients. For this purpose, we leverage a recently developed seed-inference algorithm for Alzheimer’s pathology and tailor it to the parkinsonian context. Torok et al., 2018 combined a network diffusion model that successfully recapitulates patterns of regional brain atrophy (Raj et al., 2012) with an L1-penalized optimization routine to infer the origins of pathology across individual subjects from the Alzheimer’s Disease Neuroimaging Initiative (ADNI) public database. The key requirement of this seed-inference algorithm is a *forward* model, which can predict, from a given patient’s baseline seed pattern their ongoing progression of longitudinal atrophy. Since we have already shown that the spatiotemporal dynamics of PD pathology spread can be given quantitatively and deterministically from the NDM (Pandya *et al.*, 2018), we chose it as the forward model. Following Torok *et al.*, 2018, we then design an inference algorithm which minimizes a cost function involving the forward model, and an additional penalty term that penalizes non-sparse solutions. Hence, our algorithm optimizes two constraints: (i) the *seed pattern*, *when extrapolated to future time*, must match the patient’s observed regional atrophy values, and (ii) the *seed pattern must be sparse*, such that only a small set of incipient regions will exhibit early vulnerability.

We apply our seed inference algorithm to a large public study of PD patients called the Parkinson’s Progressive Marker Initiative (PPMI) (www.ppmi-info.org/data). The same cohort was used in our previous NDM study (Pandya *et al.*, 2018), with 232 patients diagnosed with PD along with 117 age-matched controls (see **Table 1**). On this data we demonstrate that the group average pattern of inferred seeding is in strong agreement with known early sites in the brainstem, striatum and limbic areas. Intriguingly, however, there is considerable individual variability in seeding, suggesting that a common incipient state of neurodegeneration cannot explain the intersubject variability observed empirically. Thus, our methods demonstrate that the early stages of the disease may exhibit seeding patterns that are more complex and variable than generally appreciated. We identified several potential prodromal seeding variants based on clustering analysis, which, if verified in future studies, could reveal new insights into the sources of etiologic variability of PD.

**Table 1.**
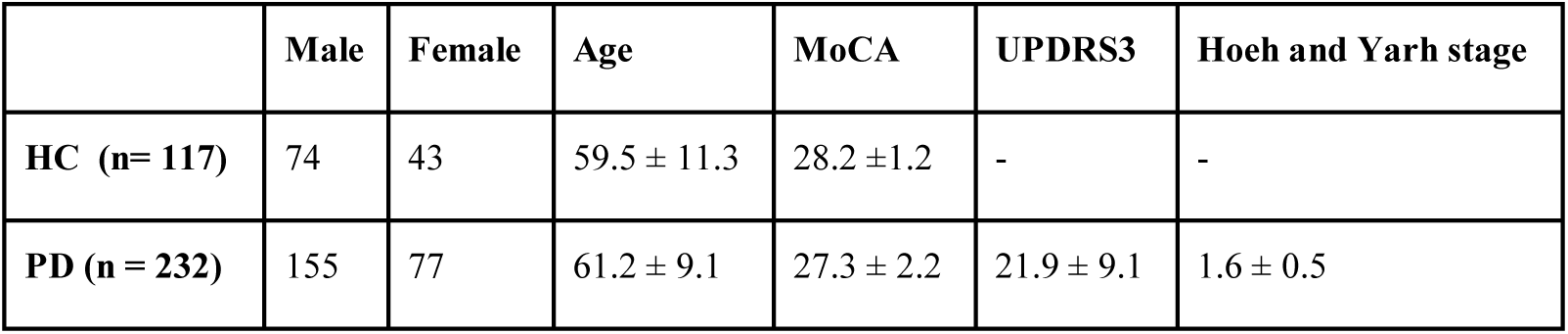
Demographic and clinical details of PPMI cohort. MoCA = Montreal Cognitive Assessment, UPDRS = Uniform Parkinson’s disease Rating Score

## 2. Materials and Methods

### 2.1 Participants and longitudinal MRI morphometric processing

A cross-sectional imaging dataset from PPMI was used to test our model for current study. 3T high-resolution T1-weighted MRI images from the baseline visit of 232 de-novo Parkinson’s patients as well as 117 age matched Healthy controls (HC) were obtained from PPMI database to measure PD related regional atrophy. Demographic and clinical features of our study are show in **Table 1** at the time of the study. The images were denoised, normalized, and corrected for non-uniformity intensity. All images were first linearly and then non-linearly registered to the MNI-ICBM 152 template. Regional brain atrophy was computed using deformation-based morphometry (DBM) (i.e. the determinant of the Jacobian transformation matrix obtained from nonlinear transformation fields). We extracted mean regional DBM values per subjects after calculating voxel-wise DBM maps. A two tailed t-test was performed between PD and HC mean DBM values to calculate the effect size of PD related regional atrophy which is represented by a vector (N=112). This t-statistic was converted to the natural range [0,1] using the logistic transform, following (Raj *et al.*, 2015). These atrophy measures were then used to test the modeling analyses.

For this study, we used a brain parcellation with 112 regions whose details have been reported previously (Zeighami *et al.*, 2015). Thirty-four cerebellar regions were removed, leaving 78 cerebral regions. In brief, supratentorial regions i.e. cortical and basal ganglia related regions were included from Hammers atlas (Hammers *et al.*, 2003). Furthermore, three midbrain structures, subthalamic nucleus, substantia nigra, and red nucleus were manually segmented using the high-resolution MRI template (T1-weighted ICBM152 template, resolution = 0.5 mm^3^), the BigBrain (Amunts *et al.*, 2013), and the brainstem anatomical atlas of (Duvernoy, 1995). A subcortical atlas based on ultrahigh-field MRI (Keuken *et al.*, 2014) was used to confirm the accuracy of the segmentations.

### 2.2 White matter connectome

An Illinois Institute of Technology Human Brain atlas v.3 constructed from high-resolution diffusion weighted MRI (DW-MRI) data from 72 young healthy subjects was used for structural connectivity. We used the anatomical connection density (ACD) as the measure of connectivity for this paper which is defined as the fraction of the connected superficial nodes with respect to the total number of superficial nodes of both areas, as proposed in (Iturria-Medina *et al.*, 2007). ACD accounts for correcting varying brain region sizes as it is obtained by dividing the raw connection strength value by the sum of region-pair surface areas. We refer to this network using the *connectivity matrix C* = {*c*_*r*,*j*_} whose elements *c*_*i*,*j*_ represent the connection strength of white matter fiber pathways between *i*^th^ and *j^th^* gray matter regions. Connections are assumed to be bidirectional due to limitations of the DTI tractography data.

### 2.3 Predictive PD Network Diffusion Model

Raj *et al.* (2012) demonstrated that the spread of proteinopathic agents over time is well captured by a dynamical system defined over a network-graph rendering of the brain, with the nodes representing gray matter structures and inter-regional connections defined as above. The *forward* Network Diffusion Model (NDM) for the pathology load **x** at time t is given by

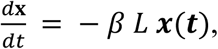

where L is the Graph Laplacian matrix given by

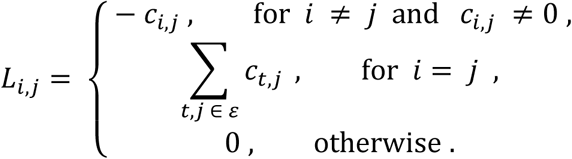

The NDM has a closed form analytical solution,

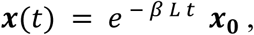

where ***x*_0_** = *x* (*t* = 0) is referred to as the seed vector or incipient atrophy pattern on which the diffusion kernel *e*^−*βLf*^ acts as a temporal and spatial blurring operator for the connectivity matrix C. In general, the unit of model’s diffusion time *t* is arbitrary and global diffusivity *β* is unknown. We chose a value that would roughly span Parkinson’s progression (10-20 years), giving *β* = 0.15. Thus, for a given initial atrophy configuration, we can use the NDM to predict the atrophy pattern at all future time points. We have previously validated the accuracy of the NDM predictions in multiple contexts (Raj *et al.*, 2012, 2015; Pandya *et al.*, 2017; Raj and Powell, 2018).

**Glossary of symbols**
| Symbol | Description |
| --- | --- |
| $\mathbf{b}$ | Empirically-observed atrophy vector |
| $\langle \mathbf{b} \rangle_{PD}$ | Average of $\mathbf{b}$ across all PD patients |
| $\mathbf{x}_{seed}$ | Seed vector, incipient atrophy pattern |
| $\langle \mathbf{x}_{seed} \rangle_{PD}$ | Average of $\mathbf{x}_{seed}$ across all PD patients |
| $\mathbf{a}(t, \mathbf{x}_{seed})$ | Predicted atrophy vector via NDM |

### 2.4 Seed-inference method

Here we recapitulate some of the key points of the seed inference method from Torok *et al.*, 2018. The empirical atrophy z-scores are given by

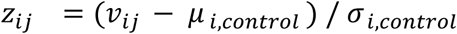

where the indexes denote the *i*-th brain region and *j*-th patient respectively; *μ* and *σ* denote mean and standard deviation calculated with respect to the healthy controls. The scores were then normalized by a weighted logistic transform to keep values within the (0,1) range; these normalized vectors will be referred to in what follows as *empirically*-*observed atrophy vectors* **b** against which we run our inference algorithm (Raj *et al.*, 2012; Torok *et al.*, 2018).

The forward NDM can be used to infer the most likely pattern of disease seeding **x**_seed_ from a given vector **b**. This inverse seed-inference process utilizes a constrained optimization algorithm with a *L*_1_-penalized cost-function to promote sparsity while maximizing the Pearson correlation R between the NDM-predicted atrophy vector **a**(*t*,*χ_seed_*) and **b**.

Analogous to the original formulation of the NDM, we define the two-variable function **a** as:

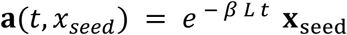

Following Torok *et al.*, 2018, we perform the minimization over the two active variables, t and **x**_seed_, in two steps. After determining an initial guess for the seed pattern, 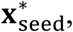 and running the NDM on each region individually, we find an estimate for the *t_min_* that satisfies the following condition:

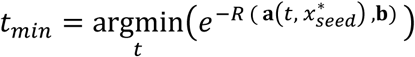

where 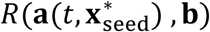 is the Pearson correlation between the NDM-predicted atrophy using initial atrophy 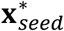 and the *empirically*-*observed atrophy vector* **b**. This cost function is monotonically decreasing and is minimized when the Pearson correlation is maximized, which is consistent with the NDM criterion. We then use this estimate of *t_min_* to find the **x**_seed_ that minimizes the following *L*_1_-constrained cost function:

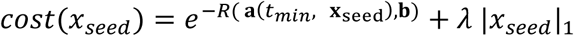

*λ* is a tunable parameter that controls the sparsity of solutions; higher values of *λ* force more entries of the resultant **x**_seed_ vector to 0. We used the same stopping criteria as Torok *et al.*, 2018, and convergence under these conditions was not an issue in all runs we observed. See the original publication for further algorithmic details.

## 3. Results

For every subject in our study, we infer an individualized seed vector **x**_seed_ ∈ ℝ^78^ that minimizes the cost function above. Each entry of the seed vector is associated with a specific gray matter region given by the brain atlas, and represents the likelihood that proteinopathic agents are present at that region when *t* = 0. When projected forward in time by the NDM, the sparse seed vectors yield predicted atrophy vectors ***a***(*t*, *χ_seed_*) that are highly correlated with their corresponding empirically-observed atrophy vectors **b**. The entries of vectors **a** and **b** will be referred to as predicted and observed atrophy-values respectively. Finally, we will use brackets and subscripts of the form < argument > _subgroup_ to indicate the average of vectors across subjects from a given subgroup (i.e. HC and PD).

## 3.1 Group-level regional patterns for atrophy and seed vectors

**Figures 1A-C** show the axial, coronal, and sagittal glassbrain views respectively of < **b** > _*PD*_, i.e., the average of the empirically-observed atrophy vectors across all PD patients. **Figures 1D-F** show the average of the inferred seed vectors < **x**_seed_ > *_PD_* across all PD patients. We note that < **x**_seed_ > *_PD_* differs from the seed vector inferred directly from < **b** > *_PD_*, since the averaging and seed-inferencing mathematical operations do not commute. In all panels, the load of proteinopathic agents in a brain region is proportional to the sphere diameter placed there, although actual values were scaled for improving visualization. **Table 2** compares the top-10 brain regions associated with the largest entries of < **b** > *_PD_* with those of < **x**_seed_ > *_PD_*. There is significant overlap between the top-10 regions for both vectors, especially regarding the Putamen, Pallidum, and Red Nucleus.

**Fig. 1:**
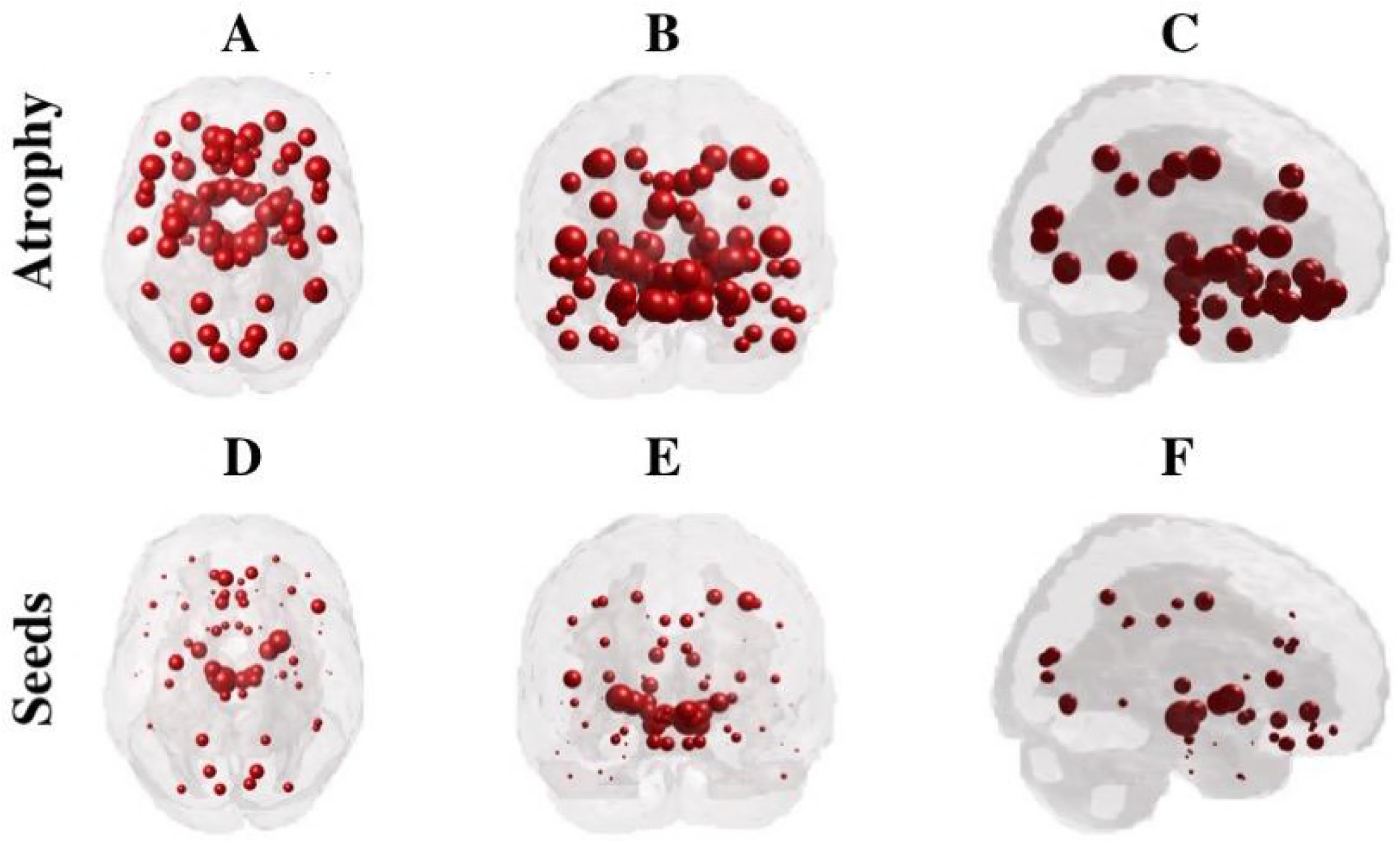
Visualization of average atrophy vector and inferred seed vector for PD patients. Panels show **(A)** the axial, **(B)** the coronal, and **(C)** the sagittal glassbrain views respectively of the average of the empirically-observed atrophy vectors across all PD patients < **b** > *_PD_*. Panels **(D-F)** show the same views for the average seed vectors across all PD patients < **x**_seed_ > *_PD_*. The pathology load in a brain region is proportional the sphere diameter placed on it. Actual values of atrophy and seed were scaled for improved visualization

**Table 2.**
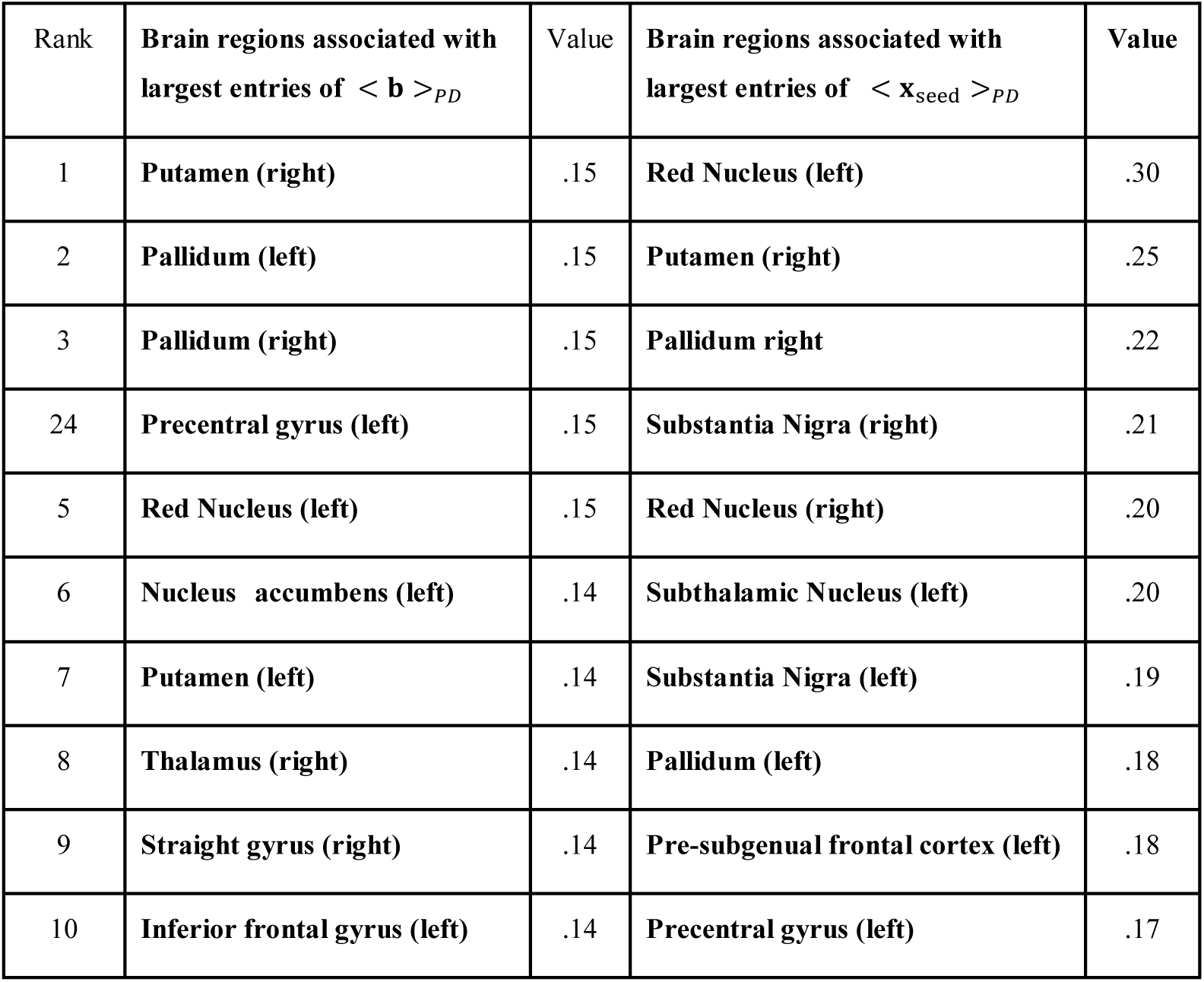
(Left) Brain regions associated with the largest entries of the empirically-observed atrophy vector averaged across all PD patients < **b** > *_PD_*. (Right) Brain regions associated with the largest entries of the inferred seed vector averaged across all PD patients < **x**_seed_ > *_PD_*

## 3.2 L-curves and parameter choices

Torok *et al.*, 2018 presented a novel method to determine a consistent seeding pattern for each patient. Naturally, the inverse-problem algorithm requires a sensible choice of the *L*_1_-penalty parameter *λ*. A priori, one does not know what this sensible choice should be, thus, it is customary to explore the parameter space within a reasonable range of values (0.1: 0.05: 1) and analyze the corresponding L-curves. The goal is to select the “elbow” of the curve, i.e., a value that provides a sensible tradeoff between the mismatch of model/data – given by the *e*^−*R*(**a**(*t_min_*, **x**_seed_),**b**)^ term in the cost-function, and the sparsity of the seed, given by the *λ* |*x_seed_*|_1_

**Figure 2** shows how the mismatch between model/data decreases as we (i) increase the value of *λ*, (ii) increase the number of seeds, or (iii) increase the *L*_1_norm of the seed. We argue that *λ* = 0.25 can be considered the “elbow” for the black curves; it provides an intermediate value of 0.63 (it is higher than the min = 0.52 and lower than the max 0.74), an intermediate average number of seeds of 6 (higher than 1 and lower than 19), and an intermediate *L*_1_-norm of 1 (higher than .2 and lower than 1.5). The curves in black are average values (±1 standard deviation) of the individualized inferred seeds. The curves in red correspond to the inferred seed from the population average atrophy-pattern. The discrepancy between curves for higher lambda values indicate the steps of the algorithm do not commute, i.e., that finding seeds and calculating their average (black) will lead to results that are significantly different than averaging the atrophies and inferring a seed from that averaged value. This strongly advocates for seed inference using patient-specific atrophy patterns as opposed to population averages, although the two curves are not too far apart for our selected value of *λ* = 0.25. All inferred seeds presented in our plots and tables were found using this value.

**Fig. 2:**
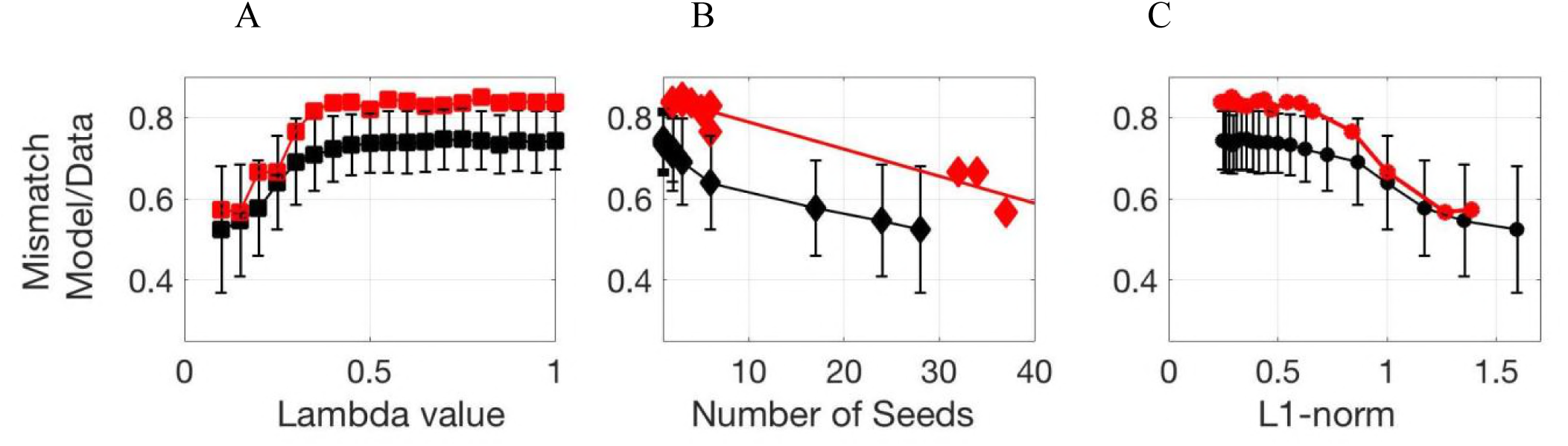
L-curves for parameter calibration. The rationale behind our choice for lambda is to make a sensible tradeoff between low mismatch between model/data and a high-sparsity for the inferred seed vector. The plots above show how the mismatch (exponential term defined in the cost function) varies in terms of **(A)** the lambda value, **(B)** the number of non-zero entries in the seed vector, and **(C)** the L1-norm of the seed vector. The curves in black are average values (±1 standard deviation) of the individualized inferred seeds. The curves in red correspond to the inferred seed from the average atrophy pattern. The discrepancy between curves for higher lambda values indicate the steps of the algorithm do not commute, i.e., that finding seeds and calculating their average (black) will lead to results that are significantly different than averaging the atrophies and inferring a seed from that averaged value. Notice that an increase in sparsity is correlated with an increase in the data/model mismatch and vice-versa. Thus, we choose lambda = 0.25 as it yields both a tractable number of non-zero entries in the seeds vector (6 out of 78) and a tolerable mismatch (value = 0.63)

### 3.3 Individualized seeds are more predictive of future atrophy

The inferred **x**_seed_ vectors varied significantly from patient to patient. In this subsection, we demonstrate that inferred **x**_seed_ are more predictive of atrophy pattern than a common seed located at a single region, i.e., a seed pattern defined by a vector with only one non-zero entry placed either in the Amygdala or in the Substantia Nigra. While certain brainstem nuclei, might be better candidate seed locations of PD pathology (e.g. medulla oblongata), these regions are not accessible on MRI and therefore we have focused on seed locations corresponding to Braak stage-III and higher.

The algorithm is as follows:

1. Infer an individualized **x**_seed_ from the observed atrophy vector **b** using the *inverse model.*
2. Project each **x**_seed_ into the future using the forward model to obtain the predicted atrophy vector ***a***(*t*,*x_seed_*). Project the single-seed vector **x**_single_ the same way to obtain ***a***(*t*,*x_Single_*).
3. Obtain the Pearson’s correlation coefficient (R-max) between (i) the pair of vectors {***a***(*t*, *x_seed_*), **b**}, (ii) the pair of vectors {***a***(*t*,*x*_*single*, *Subst*. *Nigra*_), **b**), and (iii) the pair of vectors {***a***(*t*, *x*_*Single*, *Amygdala*_), **b**}, for each PD patient. Create histograms for R-max for all cases. See Figure 3.

**Fig. 3:**
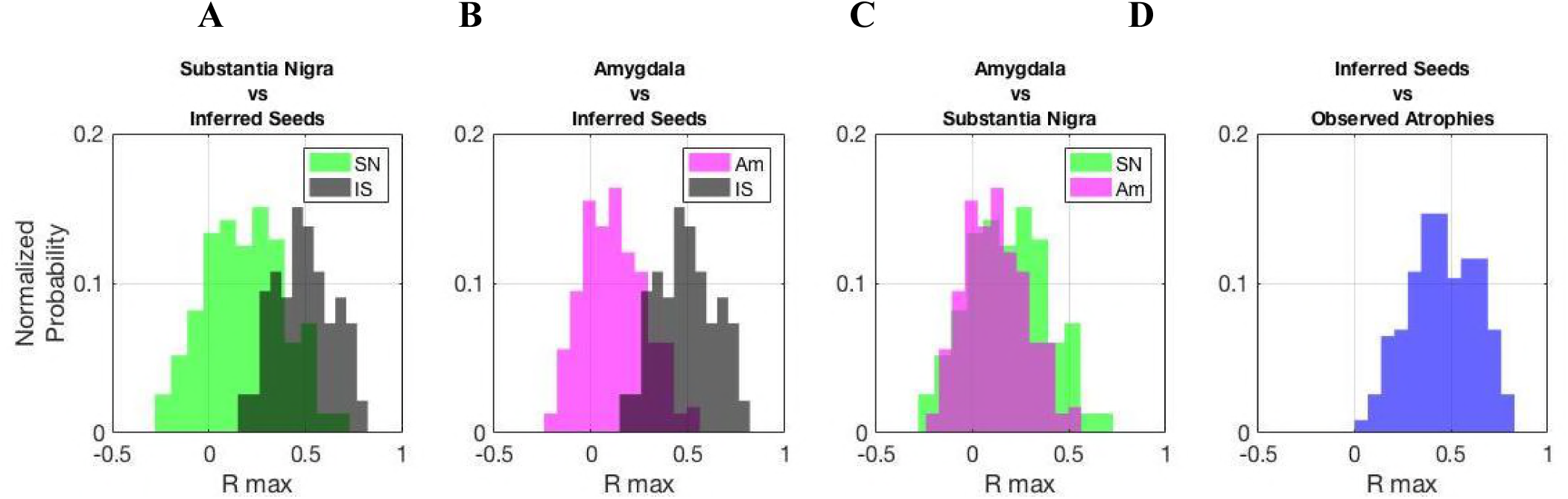
Individualized seeds are significantly more predictive of future atrophy. Our forward model estimates the progression of atrophy patterns of a given patient starting from the inferred seed pattern. Our individualized seeds lead to future atrophy patterns that are more similar to the observed ones (higher Rmax values, in black histograms) than the ones led by a single seed placed at **(A)** the substantia nigra, or **(B)** the amygdala. This result provides strong evidence against a stereotyped/standard single seed location, although **(C)** shows that a single seed in the substantia nigra is more likely than one in the amygdala. **(D)** shows that our inferred seed patterns are not obvious correlates of the observed atrophy patterns, which is consistent with the complex dynamics of disease spread in PD.

**Figures 3A-C** show the R-max histograms (i)-(iii) defined above in black, green, and in magenta respectively. The R-max values associated with (i) are typically higher than the ones associated with (ii) and (iii), demonstrating that inferred individualized **x**_seed_ patterns lead to significantly more predictive patterns of the patients’ atrophy vectors than a common single-seed vector. This result provides strong evidence against a stereotyped/standard single seed location. We also find that (**Fig. 3C**) that a single seed in the Substantia Nigra is more likely than a single seed in the Amygdala. Of course, if the seed inference algorithm was giving trivial outcomes (e.g. inferred seed pattern = observed atrophy pattern) then we would erroneously obtain similar results to the above. To guard against that possibility, In **Fig. 3D**, we present a R-max histogram comparing the two vectors (**b**, *x_seed_*}, showing that **x**_seed_ are not obvious correlates of the observed atrophy patterns. This is consistent with the complex dynamics of disease spread in PD and suggests that our seed inference is implicating a different set of regions than would be trivially predictable from the most atrophied regions.

### 3.4 Hierarchical clustering and PD subgroups

In this section we demonstrate that despite the heterogeneity of the incipient atrophy patterns across subjects, the inferred seed vectors **x**_seed_ can be categorized in two subgroups (S1 and S2). Analogously, the empirically-observed atrophy vectors **b** can also be categorized in two subgroups (A1 and A2). We use a standard technique in data mining known as standard agglomerative hierarchical cluster tree to determine, data-driven clusters in the data based on subject’s dissimilarity. In our analysis, we used MATLAB’s *linkage* function (to compute distance) with inner squared distance (Ward) and Euclidean metric.

**Figures 4A-B** summarize the sizes of all four subgroups: S1 (65 patients in cyan) vs S2 (167 patients in black) for seeds & A1 (99 patients in red) vs A2 (133 patients in blue) for atrophies. Notice that the majority of patients classified in S2 (100/167) are also classified in A2, but there is significant mixing between seed and atrophy subgroups. Visual inspection of the histograms in **Figures 4C-E** reveal that patients in all subgroups exhibit significant overlap regarding their age, and MoCA/ UPDRS3 scores.

**Fig. 4:**
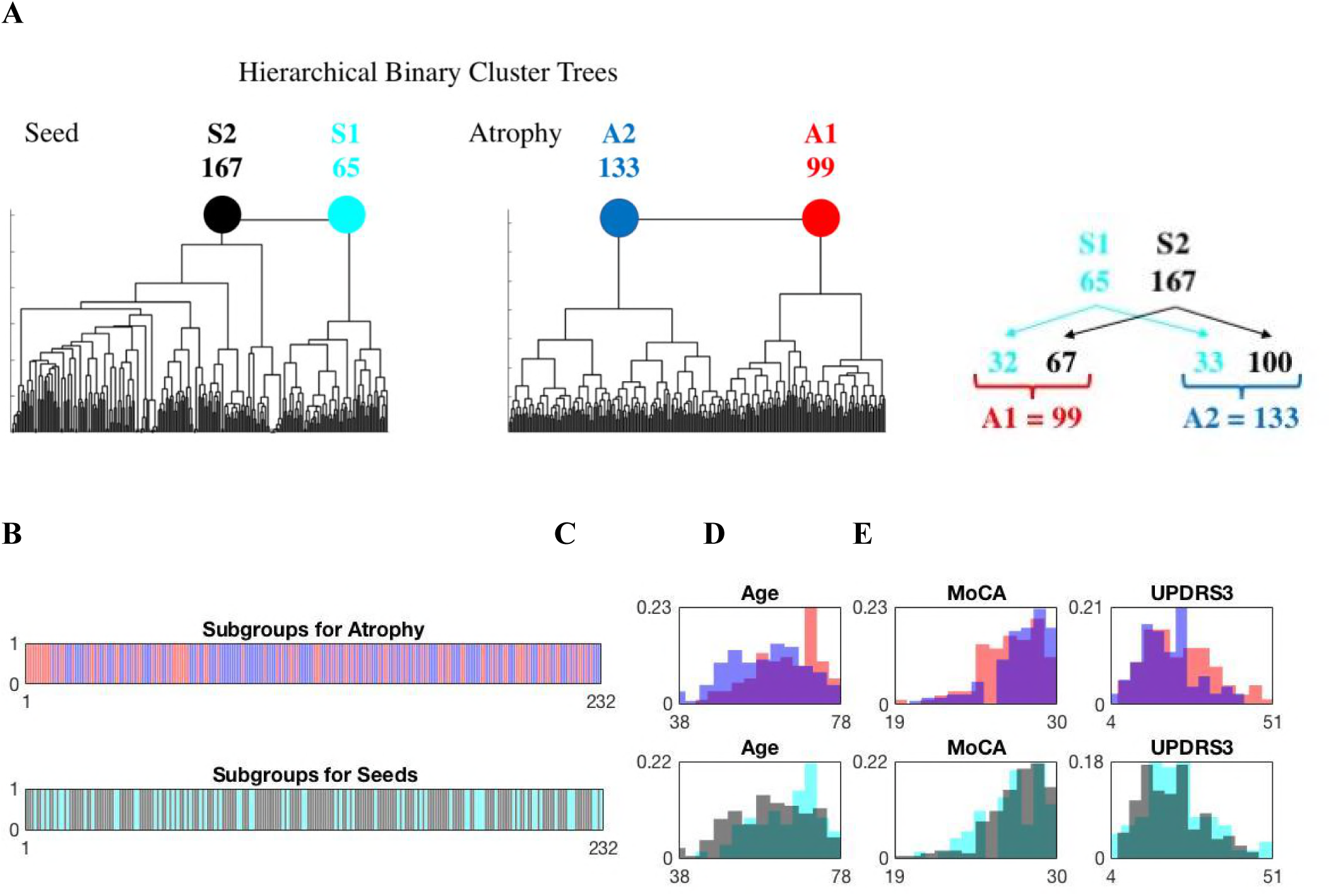
PD subgroups for seeds and atrophy. **(A)** We use MATLAB’s agglomerative hierarchical cluster tree function (linkage) to categorize the seed and atrophy data into two subgroups each: S1 (65 patients in cyan) vs S2 (167 patients in black) for seeds & A1 (99 patients in red) vs A2 (133 patients in blue) for atrophies. Distances between clusters were calculated using the inner square distance (ward) and the euclidean metric. **(B)** Classification of the 232 PD patients in our study following the color scheme above (A1 vs A2 on top, S1 vs S2 on bottom). Histograms normalized by probability for **(C)** age, **(D)** MoCA scores, and **(E)** UPDRS3 scores for each subgroup. The histograms exhibit significant overlap for all variables. Using single linkage distance (nearest neighbor) instead of the ward distance does not lead to subgroups of significant sizes as it only discriminates a few outliers. See Tables 4–5 for a detailed description of the atrophy and seed patterns for each subgroup.

**Figure 5** shows on the top of each panel, a glassbrain view of the inferred seed vectors < **x**_seed_ > *s*_1_ and < **x**_seed_ > *s*_2_ averaged between subjects in the S1 (cyan) and in the S2 (black) subgroups respectively. In the middle of each panel, we show a glassbrain view of AFS1 (cyan) and AFS2 (black). They represent the average predicted atrophy pattern (via NDM) of the seeds classified in S1 and S2 subgroups, i.e., < ***a***(*t*, *x_seed_* ∈ S1) > and <***a***(*t*,*x_seed_* ∈ S2) > respectively. Finally, in the bottom of each panel, we show a glassbrain view of < **b** > _*A*1_ and < **b** > _*A*2_, the empirically-observed atrophy vectors averaged between subjects in A1 (red) and A2 (blue) subgroups respectively.

**Fig. 5:**
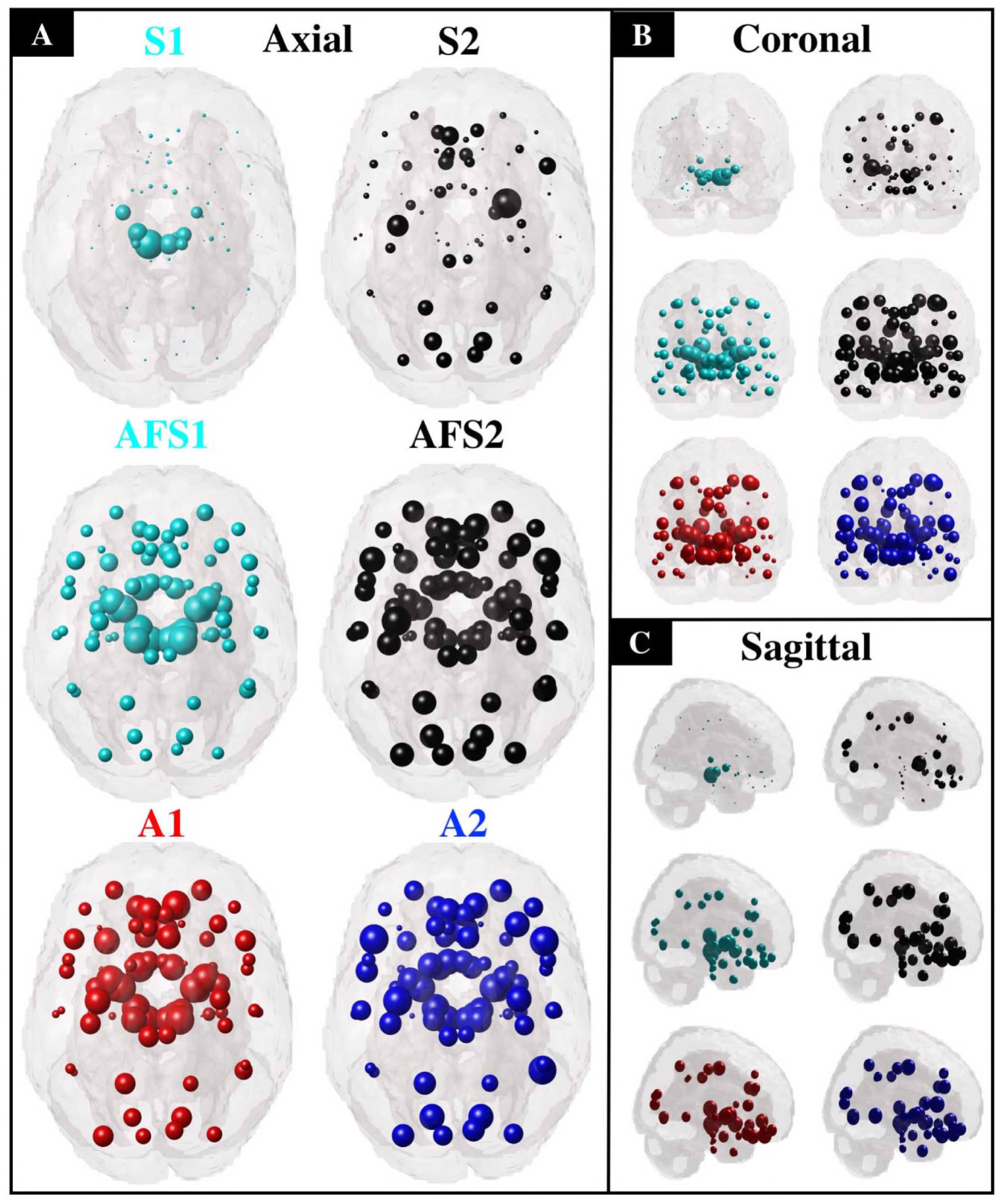
Glassbrains of the average seed pattern, projected atrophy, and observed atrophy in two subgroups found using hierarchical clustering. **(A)** Top to bottom: S1 and S2 are the individualized seed vectors averaged across subgroups S1 and S2, AFS1 and AFS2 are average projected atrophy for subgroups S1 and S2 derived from forward NDM, and A1 and A2 are observed atrophy from subgroups A1 and A2. Distinction of two subgroups from seed pattern as seen in S1 and S2 is evident as opposed to projected atrophy (AFS1, AFS2) and observed atrophy (A1, A2). We can clearly identify two different patterns, S1 showing dominant seed distribution in midbrain regions and S2 showing widespread cortical distribution. **(B)** and **(C)** shows glass brain visualization in coronal and sagittal view respectively.

**Table 3** compares the top-10 brain regions associated with the largest entries of < **x**_seed_ > _*S*1_ with those of < **x**_seed_ > _*S*2_, and **Table 4** shows the top-10 brain regions associated with those of < **b** > _*A*1_ and < **b** > _*A*2_. Finally, **Table 5** shows the regions associated with the largest entries of < ***a***(*t*, *x_seed_* ∈ S1) > and < ***a***(*t*, *x_seed_* ∈ S2) > respectively. These results are based on the hierarchical clustering analysis explained previously at two cluster level for both seed and atrophy data.

**Table 3.** The individual patient-dependent seed vectors **x**_seed_ can be categorized in two subgroups (S1 and S2) via agglomerative hierarchical cluster tree method. Left column shows the brain regions associated with the largest entries of < **x**_seed_ > _*S*1_ (in cyan) and right column shows the brain regions associated with the largest entries of < **x**_seed_ > _*S*2_ (in black).

| Rank | Brain regions associated with largest entries of $\langle \mathbf{x}_{seed} \rangle_{S1}$ | Value | Brain regions associated with largest entries of $\langle \mathbf{x}_{seed} \rangle_{S2}$ | Value |
| --- | --- | --- | --- | --- |
| 1 | Red Nucleus (left) | .49 | Putamen (right) | .34 |
| 2 | Red Nucleus (right) | .34 | Precentral gyrus (left) | .25 |
| 3 | Subthalamic Nucleus (left) | .33 | Lingual gyrus (right) | .22 |
| 4 | Substantia Nigra (left) | .31 | Pre-subgenual Frontal cortex (left) | .22 |
| 5 | Substantia Nigra (right) | .30 | Lingual gyrus (left) | .21 |
| 6 | Pallidum (left) | .26 | Inferior frontal gyrus (left) | .20 |
| 7 | Subthalamic Nucleus (right) | .26 | Cuneus (right) | .19 |
| 8 | Thalamus (right) | .20 | Superior parietal Gyrus (left) | .18 |
| 9 | Pallidum (right) | .20 | Cuneus (left) | .18 |
| 10 | Thalamus (left) | .15 | Subgenual frontal cortex (left) | .18 |

**Table 4.** The individual, empirically-observed atrophy vectors **b** can be categorized in two subgroups (A1 and A2) via agglomerative hierarchical cluster tree method. Left column shows the brain regions associated with the largest entries of < **b** > _*A*1_ (in red) and right column shows the brain regions associated with the largest entries of < **b** > _*A*2_ (in blue).

| Rank | Brain regions associated with largest entries of $\langle \mathbf{b} \rangle_{A1}$ | Value | Brain regions associated with largest entries of $\langle \mathbf{b} \rangle_{A2}$ | Value |
| --- | --- | --- | --- | --- |
| 1 | Nucleus accumbens (left) | .16 | Precentral gyrus (left) | .16 |
| 2 | Putamen (left) | .16 | Inferior frontal gyrus (right) | .16 |
| 3 | Putamen (right) | .16 | Inferior frontal gyrus (left) | .15 |
| 4 | Thalamus (right) | .16 | Posterior temporal lobe (right) | .14 |
| 5 | Pallidum (left) | .16 | Pallidum (left) | .14 |
| 6 | Pallidum (right) | .16 | Pallidum (right) | .14 |
| 7 | Medial orbital gyrus (right) | .15 | Straight gyrus (left) | .14 |
| 8 | Red Nucleus (left) | .15 | Red Nucleus (left) | .14 |
| 9 | Substantia Nigra (right) | .15 | Substantia Nigra (left) | .14 |
| 10 | Subthalamic Nucleus (left) | .15 | Substantia Nigra (right) | .14 |

**Table 5.** The individual inferred-seed vectors **x**_seed_ can be projected forward in time (via NDM) to predict the future atrophy patterns ***a***(*t*, *x_seed_*). The AFS1 (cyan) and AFS2 (black) represent the average predicted atrophy pattern of the seeds classified in S1 and S2 subgroups, i.e., < ***a***(*t*,*x_seed_* ∈ S1) > and < ***a***(*t*, *x_seed_* ∈ S2) > respectively. Left/Right column show the brain regions associated with the largest entries of each vector.

| Rank | Brain regions associated with largest entries of | Value | Brain regions associated with largest entries of | Value |
| --- | --- | --- | --- | --- |

| | $\langle \mathbf{a}(t, x_{seed} \in S1) \rangle$ | | $\langle \mathbf{a}(t, x_{seed} \in S2) \rangle$ | |
| --- | --- | --- | --- | --- |
| 1 | Thalamus (right) | .19 | Precentral gyrus (left) | .16 |
| 2 | Red Nucleus (left) | .19 | Inferior frontal gyrus (right) | .15 |
| 3 | Substantia Nigra (left) | .19 | 'Presubgen_antCing_I' | .15 |
| 4 | Substantia Nigra (right) | .19 | Nucleus accumbens (left) | .14 |
| 5 | Subthalamic Nucleus (left) | .19 | Putamen (right) | .14 |
| 6 | Anterior temporal, lobe, medial part (left) | .18 | Straight gyrus (right) | .14 |
| 7 | Pallidum (right) | .18 | Inferior frontal gyrus (left) | .14 |
| 8 | Red Nucleus (right) | .18 | Medial orbital gyrus (right) | .14 |
| 9 | Putamen (left) | .17 | Lateral remainder of occipital lobe (left) | .13 |
| 10 | Putamen (right) | .17 | Middle frontal gyrus (right) | .13 |

## 4. Discussion

The Braak staging model of PD, based on large post-mortem autopsy series, proposes that the deposition of misfolded AS occurs in a stereotypical fashion: early sites of pathological involvement are the brainstem (stages 1-2), SN and limbic structures (stages 3-4), followed by wider spread into adjoining frontal and other cortices (stages 5-6). Loss of dopaminergic neurons in the SN and other cerebral structures constitutes in an extensive neurodegenerative process (Del Tredici *et al.*, 2002; Braak *et al.*, 2003). Despite these stereotypical patterns, there is strong evidence of inter-subject variability in etiology and progression of PD patients. It is becoming increasingly evident that many patients cannot be fitted into the canonical progression scheme (Burke *et al.*, 2008; Rietdijk *et al.*, 2017). Since antemortem imaging cannot reveal directly the incipient patterns of pathology, this study aims to develop a principled, model-based method of “looking back” into the disease trajectory based on present imaging data. Many clinical uses of this approach are enumerated below, but the key outcome is to enable assessment of a patient’s incipient seeding pattern, and to use this pattern to cluster them into clinically relevant subtypes. Olfactory impairment, dementia, depression and other neuropsychiatric symptoms appear in preclinical stages of PD, preceding motor manifestations by long periods (Shiba *et al.*, 2000; Schuurman *et al.*, 2002; Leentjens *et al.*, 2003; Ross *et al.*, 2006, 2008). It is therefore plausible that the ability to infer seeding patterns of a patient can reveal likely sources of symptomatic heterogeneity.

In this paper we leverage a recently developed seed-inference algorithm for Alzheimer’s pathology (Torok et al., 2018) and tailor it to the parkinsonian context. Torok et al., 2018 combined a network-diffusion model that successfully recapitulates patterns of regional brain atrophy (Raj et al., 2012) with an L1-penalized optimization routine to infer the likely origins of pathology across individual subjects from the Alzheimer’s Disease Neuroimaging Initiative (ADNI) public database. Their results showed that the high degree of variability between patients at baseline translates to even more heterogeneous seed patterns that are significantly more predictive of future atrophy than a single seed placed in the hippocampus. Given that successful graph-theoretic models are also available for Parkinson’s Disease (Zhou *et al.*, 2012; Zeighami *et al.*, 2015; Yau *et al.*, 2018), it is sensible to investigate if similar methods are capable of revealing seeding heterogeneity in PD. In this study we found that the observed heterogeneity in PD atrophy is best explained by the heterogeneous seeding patterns. Furthermore, these seeding patterns are clustered into two main sub-groups which are not obvious form their observed atrophy. We also confirmed that inferred seeding patterns are more predictive of future atrophy patterns than a single seed placed at commonly-found sites of neuronal loss, further cementing the role of etiologic heterogeneity in PD patients. Below we discuss each of these key results in context and describe their implications.

### Group atrophy patterns of PPMI cohort

**Figure 1** (top) shows glassbrain views for the empirically-observed atrophy vectors averaged across all PD patients < b > _*PD*_. As shown in **Figures 1** and in **Table 2**, the region with the highest atrophy value is the Putamen; its primary role is to regulate movement at various stages. It plays an important role in PD because its inputs and outputs are interconnected to the Substantia nigra and Pallidus (DeLong and Wichmann, 2007). The pallidum is the next region with more atrophy.

### Group average inferred seeding patterns implicate common sites of early synuclein pathology and Braak staging

**Figure 1** (bottom) shows the average of the inferred seed vectors < **x**_seed_ > _*PD*_ from their atrophy profiles. The main regions implicated as the most likely early seeding sites in **Table 1** (SN, RN, and striatal areas like putamen and accumbens) are all regions where early PD pathology is observed (Braak *et al.*, 2003; Huot and Parent, 2007; Hanganu *et al.*, 2014; Lewis *et al.*, 2016) They also highlight a prominent role for the red nucleus in the rostral midbrain as potential *source* of the disease. It exhibits a higher seed value than regions commonly associated to PD such as the Putamen, Pallidum and Substantia Nigra. This is consistent with the notion that PD does not necessarily being in the SN (Del Tredici *et al.*, 2002; Braak *et al.*, 2003; Lang and Obeso, 2004; Ahlskog, 2005), but could also be due to the difficulty of disambiguating fiber projections to and from SN versus those that terminate at RN. Brain regions such as the Thalamus and the Nucleus accumbens were not among the top-10 seed locations despite their high atrophy values. Instead, larger seed values were found in the right Red nucleus and in the (left) Pre-subgenual Frontal Cortex.

### Inferred seeds are not simply a reflection of observed atrophy

As shown in **Table 2** and 3, the inferred group average seed regions are not the same as those with the highest levels of atrophy. This confirms that early site of PD progression are not always the areas that experience the most atrophy. It also confirms that the proposed algorithm is not trivially capturing observed atrophy, but is in fact imposing additional criteria, involving seeding sparsity and the condition that the ongoing progression occur on the anatomic network.

### Involvement of the striatum in PD seeding

The group average inferred seeding (**Figure 1, Table 1**) gives prominence to striatal structures such as putamen and accumbens. Although synuclein deposits in Lewy bodies are not quite as common in the striatum as in SNpc and the brainstem, these regions are some of the most highly affected in PD and related Lewy pathologies (Huot and Parent, 2007; Hanganu *et al.*, 2014) The fact that our inverse NDM approach was successfully able to recapitulate the key early role of these regions at the group level both confirms these regions’ prominent role in disease progression, and also provides a level of validation to the proposed inverse NDM algorithm itself. This is important, since it is currently not possible to validate our results on individual patients’ seeding patterns, as these data are in vivo, with no neuropathological confirmation or postmortem data.

Since both the empirical atrophy and our inferred seeding patterns suggest a strong striatal involvement, hence an important question is, ***why is measured striatal Lewy pathology so low?*** In PD and dementia with Lewy bodies (DLB) there has been only limited examination of striatal pathology (Jellinger and Attems, 2006). There are several possible explanations. Recent studies reveal significant striatal involvement in synucleopathies as follow. The original Braak staging of PD did not report Lewy pathology in the neostriatum (Braak *et al.*, 2003), but in a follow up they reported the presence of neostriatal lesion in stage VI of PD (Braak *et al.*, 2006). Mild striatal AS burden was associated with Braak stage 3, and very mild striatal AS lesions were seen in PD brains scoring Braak stages 3–5. Of PD with dementia (PDD) cases, 29% had positive synuclein lesions in the striatum. Mild to moderate synuclein pathology in caudate nucleus and putamen was seen in 76.5% of DLB brains. Almost 60% of DLB cases with initial PD phenotype (mean PD stage, 5.2) had some striatal synuclein lesions than PDD with mean PD stage, 4.0 (Jellinger and Attems, 2006). Numerous Lewy inclusions were reported in the putamen and neostriatum in DLB patients (Saito *et al.*, 2003). A strong correlation was found between PD stages and neostriatal inclusions (Mori *et al.*, 2008), who further reported that AS accumulates in the neostriatum at stage III initially. It is also possible that oligomeric or soluble synuclein might be more abundant than Lewy deposits in the striatum. Using novel monoclonal antibodies raised against altered synuclein, (Duda *et al.*, 2002) uncovered an extensive burden of synuclein pathology in the striatum of LB disorders, the highest density of striatal lesions being observed in patients with DLB or a combination of AD and DLB.

### Individualized seeding patterns are better predictors of future atrophy than single canonical seed regions

We find that a common incipient state of neurodegeneration cannot explain the intersubject variability observed empirically. As shown in **Figure 3**, a single-seed vector with only one non-zero entry placed either in the Amygdala or in the Substantia Nigra lead to predicted atrophy patterns that are poorly correlated with the observed data. Thus, our methods demonstrate that the early stages of the disease may exhibit incipient atrophy patterns that are more complex and variable than generally appreciated. This indicates a high level of etiologic heterogeneity in individual subjects, which makes it unlikely for any single brain region to be the source of PD pathology ramification in all subjects.

### Sources of etiologic heterogeneity in PD

Our most important finding is that the estimated incipient atrophy patterns exhibit a high degree of variability across subjects, perhaps more than generally appreciated (Del Tredici and Braak, 2016). The hierarchical clustering analysis revealed an interesting subgroup structure in the seeding patterns that were not obvious from the group seeding pattern of **Table 2**. In fact, we found two very distinct subgroups (S1 and S2, see **Figs. 4–5** and **Table 3**) from the individual subjects’ inferred seeding patterns. S1 is very clearly characterized by having the Red Nucleus and Substantia Nigra as their top seed locations (all values ≥ .30) while S2 have the Putamen, Precentral gyrus (left), and Lingual gyrus (all values ≥ .21).

The most distinguishable signature for the S1 subgroup was the presence of the Thalamus (right/left) and the Lingual gyrus (right/left) for S2. Contrary to the name, this region has little to do with speech, and instead, is linked to processing vision (especially related to letters), logical conditions, and encoding visual memories. Our analysis show that its role in early stages of PD might be more important than generally appreciated.

The hierarchical clustering tree was also applied to the observed atrophy patterns, but the difference between the two major subgroups (A1 and A2) was not as distinct as the difference between the subgroups for seeds. We conjecture that the patients in S1 (cyan) should have worse motor symptoms due to the seeding in Substantia Nigra, Pallidum, Subthalamic Nucleus and Thalamus while patients in S2 would have difficulty in attention due to seeding involvement in lingual gyrus and frontal cortex.

In summary, we report a clear distinction in seeding patterns between the two subgroups S1 and S2 – one characterized by predominant involvement of brainstem and ventral nuclei, and the other by more widespread frontal and striatal cortices. See **Tables 3–5**. This might be indicative of two distinct etiological mechanisms operative in PD. To our knowledge, this is the first time such a clear brain related sub-structure has been shown in the PD cohort. Importantly, this distinction is not apparent in the individual subjects’ atrophy patterns, which do not give a clear separation between these subgroups (labeled AFS1 and AFS2 in Figure 6), nor from a separate clustering directly from atrophy (labeled A1 and A2 in Figure 6). This suggests that while incipient seeding patterns show heterogeneity and clear sub-groups, these distinctions might be lost by the time the patients’ atrophy patterns were measured. We believe these data point to a potentially new and unreported process of progressive convergence of PD patients, starting from divergent seeding patterns.

**Fig. 6:**
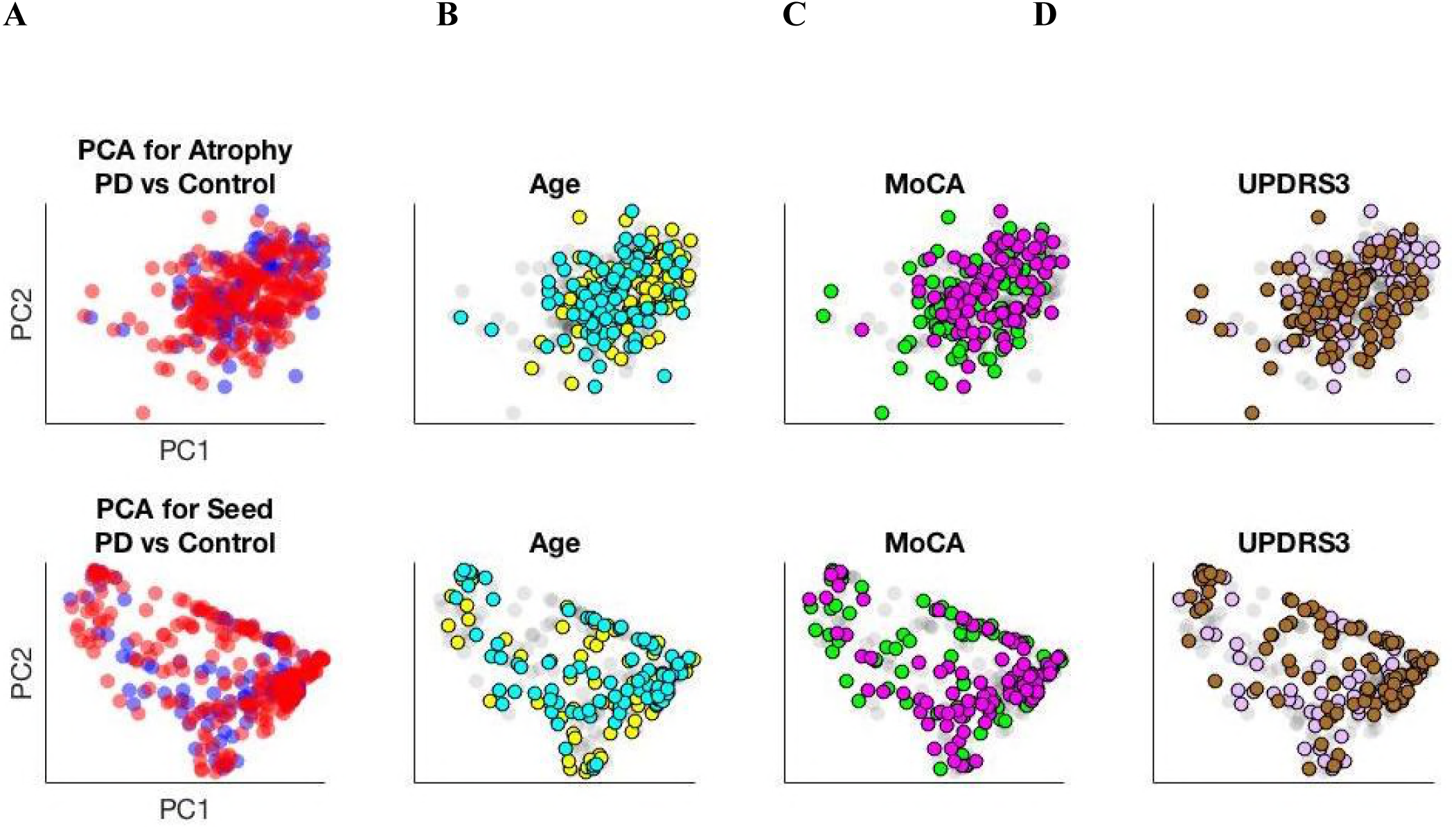
PCA for atrophy and sed patterns. Principal Component Analysis (PCA) for **(A)** atrophy and seed patterns. We form the full-data atrophy matrix X_atrophy_ (for plots on top) and the full-data seed matrix X_seed_ (for plots on bottom) from the 117 healthy controls (in blue) and the 232 PD patients (in red). The SVD decomposition allows us to project the data along the two first principal components (PC1 and PC2, that varies from top to bottom). We use distinct colors to mark in the mixed population the top/bottom **(B)** age quartiles (cyan/yellow), **(C)** MoCA scores (magenta/green), and **(D)** UPDRS3 scores (brown/beige). While there seems to be less overlap between the extreme quartiles in the bottom plots, there are no distinct clusters in any of them, which shows that the high variability of atrophy patterns within the different subjects translate also into heterogeneous seed patterns.

### The challenges of validation

While we successfully infer likely sites of disease initiation from a numerical and computational point of view, our results inherit other limitations from such as the lack of a ‘gold standard’ for neuropathological validation since our analysis are based on live patients at various stages of post-onset progression (Torok *et al.*, 2018). We instead attempt to demonstrate whether the seeds are plausible based on: (i) agreement with known pathology trends in early PD; (ii) how well they are able to predict the atrophy pattern; and (iii) if they contain latent substructure that agrees with other neuropathological markers.

The regions with the highest seed values in **Table 2**, largely located within the midbrain, show strong agreement with accepted structures that are affected earlier rather than later in AS pathology in PD (Braak *et al.*, 2003). While our seeds do not contain the more caudal structures implicated in the earliest stages of the Braak scheme, there is disagreement about the extent to which detectable α-synucleinopathy in those regions is necessary or sufficient for pre-determining rostral manifestation of PD (Burke *et al.*, 2008). In any case, since significant atrophy changes are known to occur in the amygdala (Harding *et al.*, 2002) and in substructures of the basal ganglia such as the substantia nigra (Davie, 2008), we can determine if our individual inferred seeds are more predictive of future atrophy patterns than a single seed placed at these locations. We demonstrate that the NDM predicts patient atrophy patterns using individual seeds better than a consensus seed placed either at the SN or the amygdala (**Figure 3**), similar to the results in Torok et al., 2018 for AD; in that study, it was also demonstrated that using a consensus seed containing 5 or 10 regions also resulted in poor predictions, indicating that this effect is independent of seed sparsity. In contrast to that study, however, we did detect two distinct subpopulations of patients with our seeds, one of which indicated strong midbrain involvement and the other more cortical area involvement (**Figure 5, Table 3**). The latter subpopulation is surprising in light of the generally accepted notion that the cortex harbors AS pathology only late in disease, but we note that the resultant predictions of atrophy under the NDM from the seeds of these two divergent populations are remarkably similar (R = 0.55 for AFS1 vs AFS2 in contrast to R = −0.12 for S1 vs S2). The complicated relationship between atrophy, AS pathology, and other clinical markers of PD, which limits the interpretation of these two seed subpopulations, is discussed further below

### Other limitations

It is important to remark that both incipient/observed atrophy patterns are still a long way off from more tangible clinical markers such as the Montreal Cognitive Assessment (MoCA) or the Unified Parkinson’s Disease Rating Scale 3 (UPDR3). In fact, **Figure 6** shows a Principal Component Analysis (PCA) for both atrophy and seed patterns. We use distinct colors to mark in the mixed population the top/bottom (B) age quartiles (cyan/yellow), (C) MoCA scores (magenta/green), and (D) UPDRS3 scores (brown/beige). While there seems to be less overlap between the extreme quartiles in the bottom plots, there are no distinct clusters in any of them, which shows that the high variability of atrophy patterns within the different subjects translate also into heterogeneous seed patterns. **Figure 7** shows analogous plots for data matrices restricted to PD patients alone (excluding controls). As a consequence, it remains a challenge to relate discrepancies in regional brain volumes to cognitive dysfunctions, in particular to all the different domains affected during PD such as attention and concentration, executive functions, memory, language, visuo-constructional skills, conceptual thinking, calculations, and orientation.

**Fig. 7:**
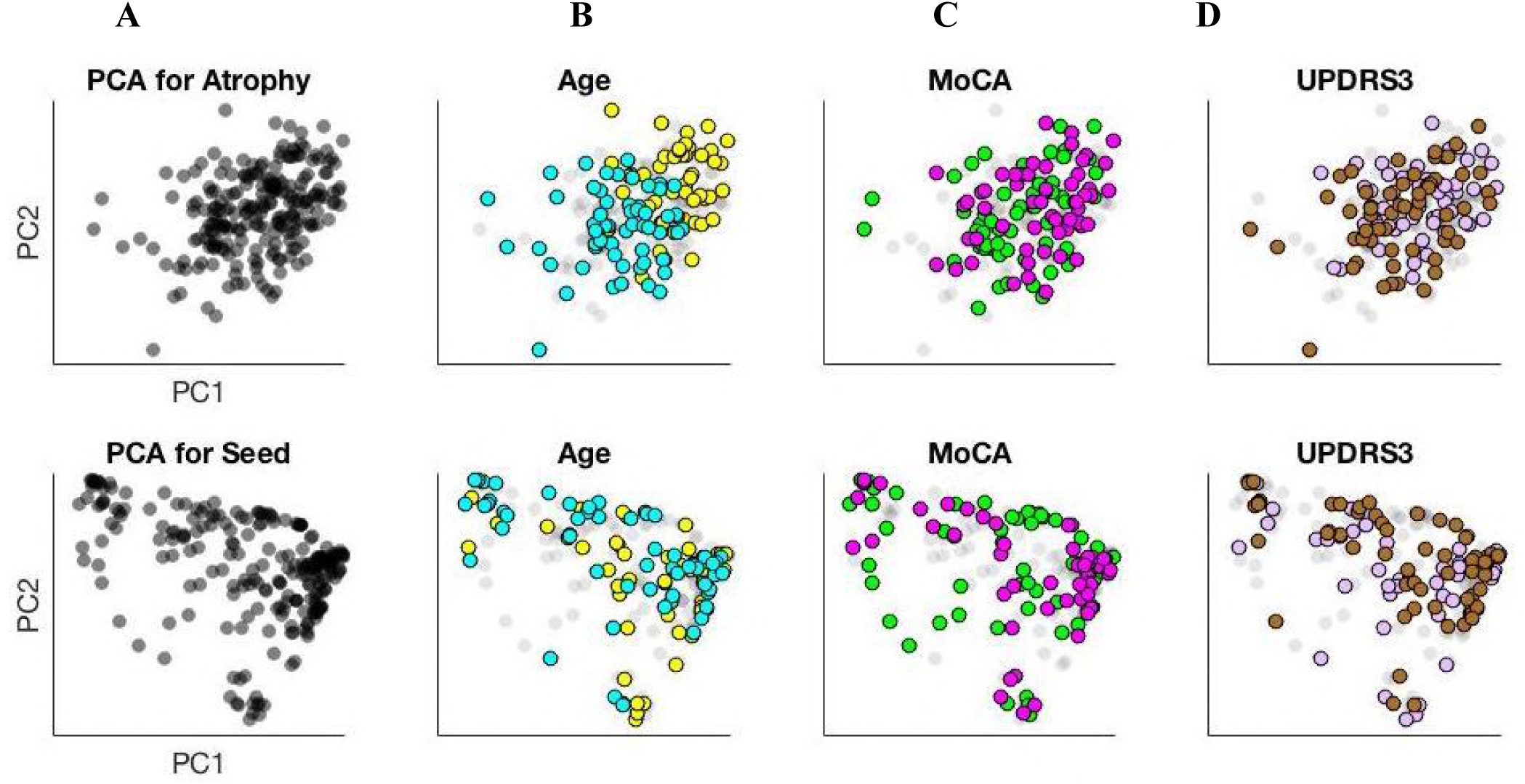
PCA for atrophy and seed patterns. Principal Component Analysis (PCA) for **(A)** atrophy and seed patterns. We form the atrophy matrix X_atrophy_ (for plots on top) and the seed matrix X_seed_ (for plots on bottom) restricted to PD patients alone (excluding controls). The SVD decomposition allows us to project the data along the two first principal components (PC1 and PC2, that varies from top to bottom). We use distinct colors to mark in the mixed population the top/bottom **(B)** age quartiles (cyan/yellow), **(C)** MoCA scores (magenta/green), and **(D)** UPDRS3 scores (brown/beige). While there seems to be less overlap between the extreme quartiles in the bottom plots, there are no distinct clusters in any of them, which shows the high variability of atrophy patterns within the different subjects translate also into heterogeneous seed patterns.

## Acknowledgements

The authors would like to thank C. Mezias and other members of the IDEAL lab for their insightful comments. AR and PM were supported by the NIH grant R01 EB022717. AR, SP, AG and JT were supported by the NIH grant R01NS092802.

## Abbreviations

PD: Parkinson’s Disease
SN: Substantia Nigra pars Compacta
PPMI: Parkinson’s Progressive Marker Initiative
NDM: Network Diffusion Model

